# A pomegranate peel extract as inhibitor of SARS-CoV-2 Spike binding to human ACE2 (*in vitro*): a promising source of novel antiviral drugs

**DOI:** 10.1101/2020.12.01.406116

**Authors:** Annalisa Tito, Antonio Colantuono, Luciano Pirone, Emilia Pedone, Daniela Intartaglia, Giuliana Giamundo, Ivan Conte, Paola Vitaglione, Fabio Apone

## Abstract

Plant extracts are rich in bioactive compounds, such as polyphenols, sesquiterpenes and triterpenes, with potential antiviral activities. As the dramatic outbreak of the pandemic COVID-19, caused by the SARS-CoV-2 virus, thousands of scientists are working tirelessly trying to understand the biology of this new virus and the disease pathophysiology, with the main goal to discover effective preventive treatments and therapeutic agents. Plant-derived secondary metabolites may play key roles in preventing and counteracting the rapid spread of SARS-CoV-2 infections by inhibiting the activity of several viral proteins, in particular those involved in the virus entry into the host cells and its replication. In this study, by using different *in vitro* approaches, we uncovered the role of a pomegranate peel extract in attenuating the interaction between the SARS-CoV-2 Spike glycoprotein and the human Angiotensin-Converting Enzyme 2 (ACE2) receptor, and in inhibiting the activity of the virus 3CL protease. Although further studies will be determinant to assess the efficacy of this extract *in vivo*, our results open up new promising opportunities to employ natural extracts for the development of effective and innovative therapies in the fight against SARS-CoV-2.

## Introduction

Plants synthesize a large variety of secondary metabolites having a wide range of biological activities and vital roles for plant survival in the environment^1^. Most of those metabolites serve to the plant as defense chemicals against both biotic stresses (e.g. herbivore insects, parasitic nematodes and microbial pathogens) and abiotic stress (e.g. low or high temperatures, deficient or excessive water, high salinity, heavy metals and ultraviolet radiations)^2^. For centuries humans used plant extracts for medicinal and health beneficial purposes, even though the active compounds responsible for the extract efficacy were mostly unknown. Thousands are the examples on the use of plant derived compounds as drugs, nutraceuticals and cosmetic ingredients^3-5^. The active compounds within plant extracts are mainly secondary metabolites that can be classified into four main categories according to their different chemical properties and structures: terpenoids, polyphenols, nitrogen and sulfur containing compounds^6^.

Polyphenols are the largest and widely distributed group of bioactive compounds in the plant kingdom. They have a distinctive structural skeleton consisting of one or more aromatic phenyl rings connected to hydroxyl groups and exhibit a wide spectrum of health properties including antioxidant protection, anti-inflammatory, anti-allergic, anti-atherogenic and anti-cancer^7,8^. Moreover, several studies demonstrated the antiviral potential of some classes of polyphenols against Epstein-Barr virus^9^, enterovirus 71^10^, herpes simplex virus (HSV)^11^, influenza virus^12^, and other viruses causing respiratory tract-related infections^13^. The mechanisms underpinning the antiviral activity of polyphenols are various (for a review see Denaro et al., 2019 ^14^), including for example the inhibition of the virus entry due to their permanent attachment on the virion envelope^15^ or the inhibition of the enzyme responsible for the virus replication^16^. The Severe Acute Respiratory Syndrome Coronavirus-2 (SARS-CoV-2) is a zoonotic pathogenic virus identified for the first time in December 2019^17^, it is responsible for one of the most serious pandemics of human history, the Coronavirus Disease 19 (COVID-19): so far the number of COVID-19 cases amounts to over 60 millions of people with more than 1.4 million deaths all over the world^18^. SARS-CoV-2, as other coronaviruses, is an enveloped positive-sense single stranded RNA virus exposing a highly glycosylated Spike (S) protein on its surface, which facilitates the viral entry into host cells. Entry depends on the binding of the surface unit S1 (portion of the S protein) to the cellular receptor Angiotensin-Converting Enzyme 2 (ACE2), facilitating viral attachment to the surface of target cells^19^. Upon binding of the S protein to the host ACE2, the virus uses the cellular serine protease TMPRSS2 for the priming of S protein itself^20^. The transcription of TMPRSS2 is promoted by androgen receptors, which could explain the predominance and the severity of pathological signs in COVID-19-affected men compared to women^21,22^, the higher proportion of men’s hospitalization^23^ and their higher mortality rates^24^.

Even though recently alternative molecular mechanisms were hypothesized to explain the virus entry into the cells^25,26^, the binding of SARS-CoV-2 S protein to human ACE2 still remains the main route of virus access to the cells and more directly related to subsequent levels of infectivity^27^. After the virus entry, the RNA genome is released into the cytoplasm and translated into two polyproteins using the translational machinery of each host cell. The two polyproteins are cleaved into the virus proteins by the main protease M^pro 28^, also referred as 3CL^pro^, and the papain-like protease PL^pro 29^, while the RNA gets replicated by its own RNA dependent RNA polymerase^30^. Once the components are all assembled, matured and packaged into new viral copies, the viruses can then exit the host cell via exocytosis and continue their infection cycles. Sars-CoV-2 mainly targets the respiratory system, intestine, cardiovascular tissues, brain and kidneys because these organs have the highest expression of ACE2^31^, resulting in symptoms such as fever, headache, dry cough and dyspnea^32^. Up to now, there are no generally proven effective therapies for COVID-19 and no vaccine is available yet, even if 350 drugs and 179 vaccines are under development, among which 56 have been employed in human clinical trials^33^. As reviewed by Dube *et al*. in October 2020^34^ antivirals can be broadly categorized into two classes: the first includes those targeting viral proteins involved in viral life cycle or in virus structure, and the other mostly targets host proteins which are important for viral infection or for the host’s immune response.

A large number of plants derived compounds are under investigation for their potential therapeutic effects against SARS-CoV-2. Many reports based on molecular docking analysis suggested the potential capacity of polyphenols, such as curcumin, kaempferol, catechin, naringenin, quercetin^35^ or hesperidin, rutin and diosmin^36^ to inhibit the activity of SARS-CoV-2 main protease and consequently the virus replication. One study also suggested that the binding of two polyphenols, punicalagin (PC) and theaflavin, to S protein could be exploited as strategy to inhibit the virus entry into human cells^37^.

Pomegranate (*Punica granatum* L.) fruits, extensively produced by Mediterranean countries, including Tunisia, Turkey, Egypt, Spain, Morocco and Italy, are rich in polyphenols, such as ellagitannins (ETs), mainly including α and β isomers of punicalagin (PC), gallic acid (GA), ellagic acid (EA) and its glycosylated derivatives, and anthocyanins^38^. The pomegranates are majorly processed by food industries to obtain juices or jams from the arils, while the peels, that constitute around 50% of the fresh fruit weight, are discarded. It has been reported that the peels had a higher content of dietary fiber and total polyphenols, as well as a stronger antioxidant capacity than the pulp fraction of the fruit itself, thus they could be a valuable source of extracts for cosmetic and nutraceutical applications^39^. Several evidences suggest that these compounds may have protective activity against degenerative chronic diseases, such as some types of cancer, type 2 diabetes, atherosclerosis and cardiovascular diseases^40,41^. Furthermore, a number of studies on pomegranate peel extracts focused on their antibacterial and antiviral activity^42^ as well as on the property to inhibit influenza^43^ and Herpes virus replication^44^. These observations let hypothesize that pomegranate peel extracts may be employed as antiviral ingredients against SARS-CoV2. Therefore, the aim of this work was assessing the potential of pomegranate peel extracts to counteract SARS-CoV2 infection. We found that a hydroalcoholic extract obtained from pomegranate peels (PPE) and its main constituents were able to inhibit the binding between SARS-CoV-2 S glycoprotein and ACE2 *in vitro*, suggesting a potential of the extract in the prevention of SARS-CoV-2 entry into host cells. Moreover, PPE compounds inhibited the virus 3CL protease, suggesting a potential use of the extract as natural remedy to enhance protection against SARS-CoV-2.

## Material and methods

### Preparation of PPE

Dried pomegranate peels were provided by Giovomel, an Italian company producing pomegranate juice. The preparation of the Pomegranate Peel Extract (PPE) was performed by adding 700 mL of a solution ethanol/water (70/30, v/v) to 150 g of dried peels, at 4°C, according to Malviya *et al.*, 2014^45^. The mixture was homogenized 3 min at 1500 rpm and 2 min at 3000 rpm by using a Grindomix GM 300 knife mill (Retsch GmbH, Haan, Germany). The resulting suspension was left under stirring at 150 rpm for 2 h at 25°C, avoiding light exposure. The suspension was then centrifuged at 6300 rpm for 10 min at 4°C. The supernatant was filtered through a filter paper (FILTER-LAB, qualitative filter paper, Barcelona, Spain) and concentrated under vacuum in a rotary evaporator (IKA RV8, IKA-Werke GmbH & Co, Staufen, Germany) set to 25°C. Finally, the pH of the concentrated extract was adjusted to 7.0 with 10N NaOH and then freeze-dried until obtaining a fine powder.

### High Resolution Mass Spectrometry (HRMS) analysis of PPE

LC-MS data were acquired on an Accela U-HPLC system coupled to an Exactive Orbitrap mass spectrometer (Thermo Fisher Scientific, San Jose, CA) equipped with a heated electrospray interface (HESI). The chromatographic separation was carried out according to Colantuono et al., 2017^46^. Briefly, we used a Gemini C18-110Å column, 150 mm × 2.0 mm, 5 μm (Phenomenex, Torrance, CA) heated to 30°C and the mobile phases consisted of 0.1% formic acid water (A) and 0.1% formic acid acetonitrile (B) with a flow rate of 200 μL/min. The dry extracts were dissolved in methanol/water (50:50, v/v) and 10 μL were injected into the column. MS data acquisition was performed in negative ionization modes, in the mass range of *m/z* 100–1300. The resolving power was set to 50,000 full width at half-maximum (FWHM, m/z 200) resulting in a scan time of 1 s. The automatic gain control was used in balanced mode (1 × 10^6^ ions); maximum injection time was 100 ms. The interface parameters were the following: spray voltage 3500 kV, capillary voltage 50V, capillary temperature 275 °C, sheath gas 30 arbitrary units and auxiliary gas 15 arbitrary units.

Calibration curves were constructed in the linearity ranges of 1-50 μg/mL for PC and 0.1-5 μg/mL for EA, GA. Metabolite identification was performed by using exact mass values up to the fifth decimal digit with mass tolerance ± 5 ppm. **Table 1** reports the polyphenols identified in PPE and individual molecular formula, retention time, theoretical mass, experimental mass and error. The amount of each compound in the extract was determined by using PC and EA as reference standards for ellagitannins (ETs) and EA derivatives (EAs), respectively. Punicalin (α, β isomers), Granatin B, Causarinin, Galloyl-HHDP-hexoside, Pedunculagin I (bis-HHDP-hex), Pedunculagin II (Digalloyl-HHDP-hex) were expressed as equivalents of PC. EA hexoside, EA pentoside, EA deoxyhexoside were expressed as equivalents of EA. Total polyphenols were calculated as sum of all the compounds retrieved.

**Table 1:**
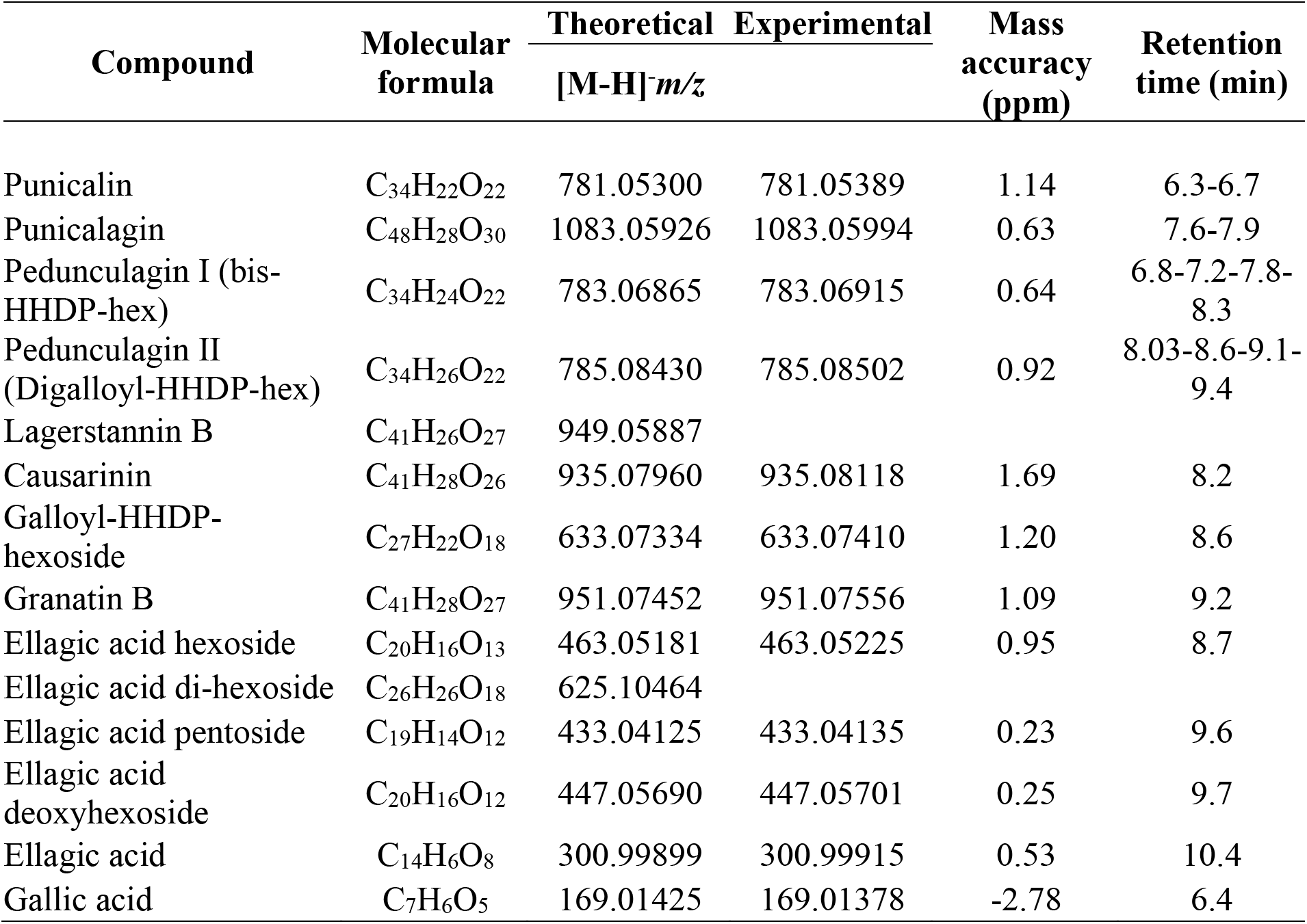
High-Resolution Mass Spectrometry identification of the compounds in PPE achieved by Orbitrap MS.

| Compound | Molecular<br>formula | Theoretical<br>[M-H]<sup>-</sup>*m/z* | Experimental | Mass<br>accuracy<br>(ppm) | Retention<br>time (min) |
| --- | --- | --- | --- | --- | --- |
| Punicalin | C<sub>34</sub>H<sub>22</sub>O<sub>22</sub> | 781.05300 | 781.05389 | 1.14 | 6.3-6.7 |
| Punicalagin | C<sub>48</sub>H<sub>28</sub>O<sub>30</sub> | 1083.05926 | 1083.05994 | 0.63 | 7.6-7.9 |
| Pedunculagin I (bis-HHDP-hex) | C<sub>34</sub>H<sub>24</sub>O<sub>22</sub> | 783.06865 | 783.06915 | 0.64 | 6.8-7.2-7.8-8.3 |
| Pedunculagin II<br>(Digalloyl-HHDP-hex) | C<sub>34</sub>H<sub>26</sub>O<sub>22</sub> | 785.08430 | 785.08502 | 0.92 | 8.03-8.6-9.1-9.4 |
| Lagerstannin B | C<sub>41</sub>H<sub>26</sub>O<sub>27</sub> | 949.05887 |  |  |  |
| Causarinin | C<sub>41</sub>H<sub>28</sub>O<sub>26</sub> | 935.07960 | 935.08118 | 1.69 | 8.2 |
| Galloyl-HHDP-hexoside | C<sub>27</sub>H<sub>22</sub>O<sub>18</sub> | 633.07334 | 633.07410 | 1.20 | 8.6 |
| Granatin B | C<sub>41</sub>H<sub>28</sub>O<sub>27</sub> | 951.07452 | 951.07556 | 1.09 | 9.2 |
| Ellagic acid hexoside | C<sub>20</sub>H<sub>16</sub>O<sub>13</sub> | 463.05181 | 463.05225 | 0.95 | 8.7 |
| Ellagic acid di-hexoside | C<sub>26</sub>H<sub>26</sub>O<sub>18</sub> | 625.10464 |  |  |  |
| Ellagic acid pentoside | C<sub>19</sub>H<sub>14</sub>O<sub>12</sub> | 433.04125 | 433.04135 | 0.23 | 9.6 |
| Ellagic acid<br>deoxyhexoside | C<sub>20</sub>H<sub>16</sub>O<sub>12</sub> | 447.05690 | 447.05701 | 0.25 | 9.7 |
| Ellagic acid | C<sub>14</sub>H<sub>6</sub>O<sub>8</sub> | 300.99899 | 300.99915 | 0.53 | 10.4 |
| Gallic acid | C<sub>7</sub>H<sub>6</sub>O<sub>5</sub> | 169.01425 | 169.01378 | -2.78 | 6.4 |

### Antioxidant activity of PPE

The antioxidant capacity (AC) of PPE was measured by using the ABTS assay as reported by Re *et al.*, 1999^47^. Briefly, a stable stock solution of ABTS·^+^ was produced by reacting a 7 mmol/L aqueous solution of ABTS with 2.45 mmol/L potassium persulfate (final concentration) and allowing the mixture to stand in the dark at 4°C for 16 h before use. The ABTS·^+^ solution was diluted with ethanol to an absorbance of 0.700 ± 0.050 at 734 nm. Freeze-dried PPE was appropriately diluted in water and 0.1 mL of reconstituted extract was added to 1 mL of ABTS·^+^ solution. The mixture was allowed to stand at room temperature for 2.5 min prior the absorbance was recorded at 734 nm by using the multiplate reader Victor Nivo (Perkin Elmer). Results were expressed as μmol Trolox equivalents (TE)/g of powder.

### SARS-CoV-2 Spike RBD/ACE2 binding inhibitor assay

The inhibition of the S-ACE2 interaction was measured using the SARS-CoV2 Inhibitor Screening Assay kit (Adipogen, Cat. N° AG-44B-0007-KI01). According to the manufacturer’s instructions, briefly 100 μl of Receptor Binding Domain (RBD) of Spike (1 μg/mL) was used for a 96 well-plate coating for 16 h at 4°C. The plate was then treated with the blocking buffer for 2 h at room temperature, washed in wash buffer and incubated with the PPE or compounds for 1 h at 37°C in the Inhibitor Mix Solution (IMS), containing biotin conjugated-ACE2 0.5 μg/mL. After incubation HRP labeled-streptavidin (1:200 dilution) was added to each well and incubated for 1h at room temperature. The reaction was developed by adding 100 μl of TMB (Tetramethylbenzidine Neogen) for 5 min at RT and measured at 450 nm by the microplate reader Victor Nivo (Perkin Elmer).

### Microscale thermophoresis

Microscale thermophoresis (MST) experiments were performed on a Monolith NT 115 system (Nano Temper Technologies, Munchen, Germany) and designed to evaluate the ability of the PPE to bind ACE2, S protein and RBD (Sino Biological, USA). The proteins used in the study were: ACE2 (NP_068576.1) (Met1-Ser740), Spike FL (YP_009724390.1) (Val16-Pro1296) and RBD Spike (YP_009724390.1) (Arg319-Phe541); all three produced as recombinant in baculovirus-insect cells and carrying a polyhistidine tag at the C-terminus. Each protein (10 μM) was labeled with NT-647-NHS reactive dye (30 μM) (Nanotemper, Germany), which reacted efficiently with the primary amines of the proteins to form a stable dye protein conjugate. PPE was used in the concentration range of 65 μM–1.92 × 10^−3^ μM in the experiment with ACE2, 32.5 μM-9.92 × 10^−4^ μM with Spike and 3.25 μM–9,92 × 10^−5^ μM with RBD Spike respectively, preparing 16-point serial dilution (1:2) in PBS supplemented with tween 0.05%. The concentration values of the extract referred to the corresponding quantity of punicalagin, the most abundant extract polyphenol, as determined by chemical analysis. The MST was carried out using 100% LED and 20% IR-laser power at 37 °C. The ligand in the experiments with Spike FL and RBD induced a quenching of fluorescence so, to confirm the specificity of interaction, the SDS denaturation test (SD-Test) was performed. An equation implemented by the software MO-S002 MO Affinity Analysis, provided by the manufacturer, was used for fitting the normalized fluorescence values at different concentrations of the ligands.

### Lentivirus infection

Human Kidney-2 cells (HK-2) were obtained from American Type Culture Collection (ATCC) and were cultured in Dulbecco’s Modified Eagle Medium (DMEM) (EuroClone, Milano Italy) supplemented with 5% (v/v) FBS, 1% Insulin-Transferrin-sodium Selenite media supplement (ITS) (Sigma-Aldrich-Merck KGaA, Germany) and 1% penicillin-streptomycin. The cells were maintained at 37°C, 5% CO2 in a humidified incubator according to the guidelines provided by the vendors, plated in 96-well plates (CellCarrier-96 ultra with lid, Perkin Elmer), at a density of 5×10^3^ per well in 100 μl culture medium. After 24 h, the cells were incubated with either 0.04 mg/mL of PPE extract or water for 4 h. The cells were then infected with SARS-CoV-2 Spike-Pseudotyped Lentivirus (Firefly Luciferase SARS-CoV-2 lentiviral particles-GeneCopoeia) and the control VSV-G protein pseudotyped Lentivirus (HLUC-Lv201 Firefly luciferase − eGFP lentifect-GeneCopoeia) at a concentration of 4,9E−9 GC/mL and 1,2E−9 GC/mL, respectively. After 72 h, the cells were fixed in 4% paraformaldehyde and washed three times in PBS. Nuclei were counterstained with DAPI and after washing the cells were imaged by the Operetta High Content Imaging System (Perkin Elmer Woodbridge, Ontario, Canada), using a 20x magnification objective. Acquired images were analyzed by the software Columbus (Perkin Elmer), version 2.6.0. Image analysis consisted of identifying and counting viral-infected HK-2 cells based on 488-intensity fluorescence. The infection rate was calculated as the ratio between the number of infected cells and the number of total cells counted per well. The plot, showing the percentage of 488-positive cells after pomegranate treatment, was compared to that in H2O-treated cells.

### Gene expression analysis on HK2 cells

Cells were plated in 24-well plates at a density of 5 × 10^4^ per well in 500 μl culture medium. After 24 h the cells were incubated with 0.04 mg/mL of PPE for 72 h and then collected for RNA extraction, performed by the GeneElute Mammalian Total RNA purification kit (Sigma Aldrich-Merck KGaA Germany). The RNA was treated with deoxyribonuclease (DNAse) I (Thermo Fisher Scientific, Dallas, TX, USA) at 37°C for 30 min. Reverse transcription was performed using the RevertAid™ First Strand cDNA Synthesis Kit (Thermo Fisher Scientific, Dallas, TX, USA). Semiquantitative RT-PCR was performed with the Quantum RNA™ kit (Thermo Fisher Scientific, Dallas, TX, USA) containing primers to amplify 18S ribosomal RNA (18S rRNA) along with competimers, that reduced the amplified 18S rRNA product within the range to be used as endogenous standard. The amplification reactions were made using specific oligonucleotides by the Mastercycler™ ProS (Eppendorf, Milano, Italy) with the following general scheme: 2 min at 94°C followed by 35 cycles of 94°C for the 30 s, 50°C for 30 s, and 72°C for 30 s, with a 10 min final extension at 72°C. The PCR products were loaded on 1.5% agarose gel, and the amplification bands were visualized and quantified with the Geliance 200 Imaging system (Perkin Elmer). The amplification band corresponding to the analyzed gene was normalized to the amplification band corresponding to the 18S and reported as a percentage of untreated controls set as 100%. The used primer sequences for the amplifications were the following: ACE2 Fw *ATGTCACTTTCTGCAGCC*; ACE2 Rv *GTTGAGCAGTGGCCTTACAT*; TMPRSS2 Fw *ATTGCCGGCACTTGTGTTCA*; TMPRSS2 Rv *ACAGTGTGCACCTCAAAGAC*.

### 5alpha Reductase activity

Hair Follicle Dermal Papilla cells (HFDPC) were seeded in a 96 well plate at a density of 8 × 10^3^, after 16 h they were stimulated with testosterone 600 nM and treated with pomegranate extract or finasteride 100 nM for 24 h. Another 96 well plate was coated with 100 ng of DHT-conjugated BSA, the day after the plate was washed with PBS − 0.05% Tween20 and incubated with a blocking solution containing PBS, Tween20 and 3% of BSA for 1 h. After 3 washes, the plate was loaded with 50 μl of cell supernatants derived from cell treatments, plus 50 μl of biotin-conjugated anti-DHT antibody (1:1000 dilution in PBS − BSA 1%). After 2h, the plate was washed 3 times and incubated with 5μg/mL of peroxidase-conjugated streptavidin for 1 hour at room temperature. After 3 washes, 0.5 mg/mL of OPD in 50 mM citrate buffer − 0.012% H_2_O_2_ was added to each well and the absorbance was measured at 490nm by the microplate reader Victor Nivo (Perkin Elmer).

### 3CL protease activity assay

To measure the activity of the viral 3CL protease in the presence of PPE extract we used the Untagged (SARS-CoV-2) Assay kit provided by BPSBioscience (CA, USA), according to the procedure described in the provider’s instructions. Briefly, 15 ng of 3CL protease was incubated with the extract at the indicated concentrations or with 500 μM of GC376, used as positive control. After 30 min of incubation at room temperature, the enzymatic reaction was carried on for 24 h by the addition of 40 μM 3CL protease substrate. The fluorescence was measured by the Victor Nivo Microplate reader (Perkin Elmer) exciting at 360 nm and detecting at 460 nm.

### Statistical analysis

All the measures were expressed as means ± standard deviations (SD) of three independent experiments. A paired-samples t-test was conducted by Microsoft Excel; a p value lower than 0.05 was considered statistically significant.

## Results and Discussion

### Chemical characterization of PPE

The concentration of polyphenols in PPE is reported in **Table 2**. ETs were the most abundant compounds. Specifically, PC represented 38.9 % of all the polyphenols detected in the extract, followed by pedunculagin anomers and punicalin anomers representing 16.7% and 13.2% of total polyphenols, respectively. These results were in accordance with previous studies published by Lu *et al.*, 2008 and Fischer *et al.* in 2011^48,49^. The sum of EAs and GA represented 3.9 % of the total polyphenols in PPE.

**Table 2:** Total amount of ellagitannins (ETs), ellagic acid derivatives (EAs) and gallic acid (GA) in PPE. The values are expressed as mg/g of dry powder (mean values ± standard deviation).

| Compounds | PPE<br>(mg/g) |
| --- | --- |
| Punicalagin | 182.31 ± 0.75 |
| Punicalin | 61.95 ± 2.34 |
| Granatin B | 61.04 ± 7.25 |
| Causarinin | 20.79 ± 2.52 |
| Galloyl-HHDP-hexoside | 45.40 ± 1.53 |
| Lagerstannin B | <LOD |
| Pedunculagin I | 50.25 ± 0.98 |
| Pedunculagin II | 28.04 ± 0.42 |
| **Total Ellagitannins** | 449.78 ± 8.31 |
| Ellagic acid | 10.71 ± 1.17 |
| Ellagic acid hexoside | 3.00 ± 0.13 |
| Ellagic acid pentoside | 1.88 ± 0.09 |
| Ellagic acid deoxyhexoside | 1.87 ± 0.11 |
| Ellagic acid diexoside | <LOD |
| **Ellagic acid derivatives** | 17.45 ± 1.49 |
| Gallic acid | 0.98 ± 0.11 |
| **Total\*** | 468.20 ± 9.69 |

\*Expressed as sum of mg of punicalagin equiv. + mg of ellagic acid equiv. + mg of gallic acid equiv.; <LOD, lower than the limit of detection.

Notably, the antioxidant capacity of PPE measured by ABTS method was 3590 μmol TE/g of extract. This value corresponded to 1041 μmol TE/g of dried pomegranate peels and was in line with the data showed by Marchi *et al.* in 2015^50^ (872-1056 μmol TE/g of dried peels) and by Fischer *et al.* in 2011^48^ (1362 μmol TE/g of dried peels and 2887 μmol TE/g of dried mesocarp).

### Effect of PPE on Spike/ACE2 binding

To assess whether PPE had an inhibitory activity on S/ACE2 binding, we used a SARS-CoV-2 inhibitor screening kit by Adipogen. PPE, used at three concentrations ranging from 0.04 mg/mL to 1 mg/mL, inhibited the interaction between S and ACE2 up to 74%, and this effect was dose dependent (**Figure 1**). As positive control, we used AC384, a monoclonal antibody that inhibited the binding between S and ACE2 by specifically recognizing ACE2 itself, accordingly to the manufacturer’s instruction.

**Figure 1:**
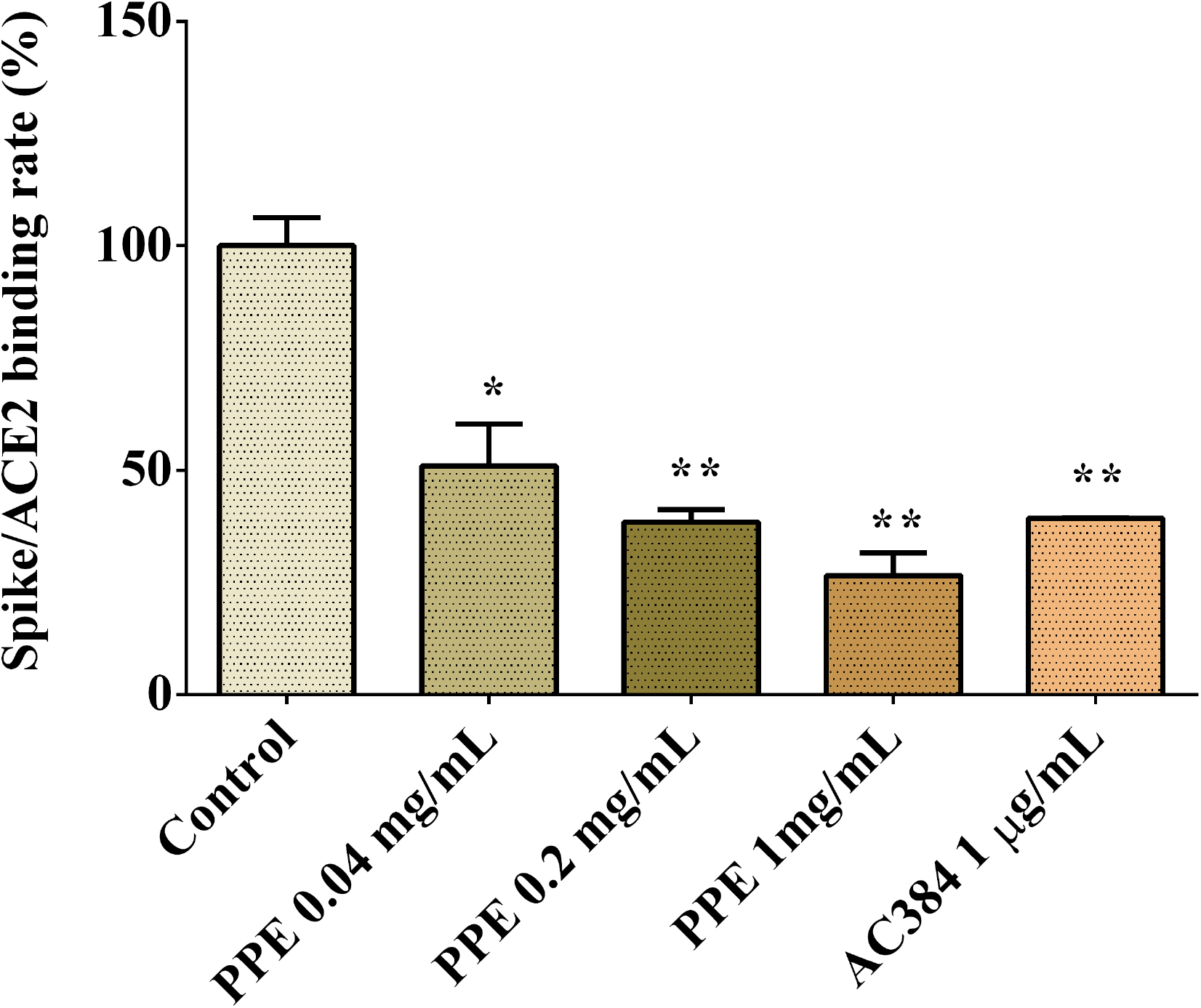
Spike/ACE2 binding in the presence of PPE, used at three concentrations compared to control and antibody inhibitor AC384. The results are the averages of three independent experiments, expressed as percentages respect to control arbitrarily set as 100%. The error bars represent standard deviations and the asterisks indicate statistically significantly values (*p value is between 0.01 to 0.05; ** 0.001 to 0.01) according to T test.

To provide insights into which of the PPE polyphenols were relevant for that inhibition, the three most abundant components of PPE, i.e. PC, EA and GA, were individually tested, at the same concentrations as present into 0.04 mg/mL PPE. The results in **Table 3** showed that PC most affected the binding between S and ACE2 by exerting 49% inhibition, followed by EA with 36% inhibition, whereas GA did not have any effect.

**Table 3:** Spike/ACE2 binding (%) in the presence of punicalagin, ellagic acid and gallic acid, at concentrations corresponding to those present in 0.04 mg/mL of PPE and equal to 7.29, 0.43 and 0.04 μg/mL, respectively. The results are the averages of three independent experiments, expressed at percentage respect to control arbitrarily set as 100%.

| Sample | Binding (%) | SD | pvalue |
| --- | --- | --- | --- |
| Spike/ACE2 | 100 | +/- 10 |  |
| Spike/ACE2 + PPE | 51 | +/- 11 | 0.04 |
| Spike/ACE2 + Punicalagin | 36 | +/- 4 | 0.01 |
| Spike/ACE2 + Gallic acid | 100 | +/- 2 | 0.5 |
| Spike/ACE2 + Ellagic acid | 64 | +/-10 | 0.03 |

To further investigate on the pomegranate compound binding capacity, the chemical interactions between the extract and S, and between the extract and ACE2, were analysed by MicroScale Thermophoresis (MST) experiments (Figure 2, S1 and S2). The results showed that the PPE bound both the proteins (**Figure 2**), even though the interaction with S was 10 folds stronger than that to ACE2. Moreover, we observed that the binding of PPE compounds to S was mostly due to a high affinity towards the Receptor Binding Domain (RBD) of the protein, as the chemical interaction to this domain was more similar to that calculated for the full-length protein.

**Figure 2:**
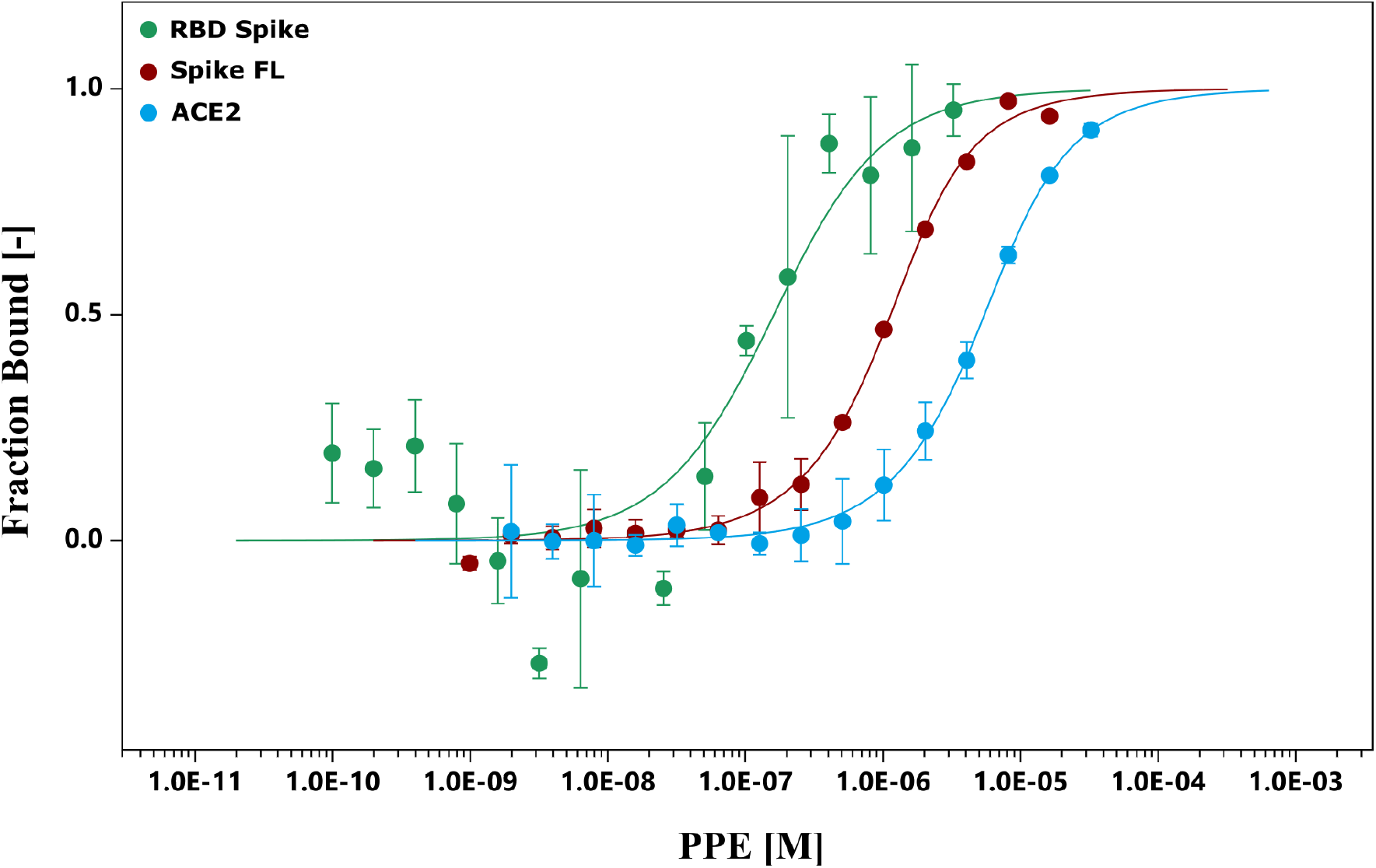
MicroScale Thermophoresis (MST). The binding curves were obtained incubating PPE with the Spike Receptor Binding Domain (RBD Spike), Spike full-length protein (Spike FL) and ACE2.

The biochemical data prompted us to investigate on the capacity of PPE to effectively inhibit the interaction between S and ACE2 in a cellular model. To do that, we used a system based on a Spike-carrying Lentivirus, infecting human renal cells (HK2), already known to express ACE2^51^. As control we used a lentivirus that did not carry S, but the Vesicular Stomatitis Virus G (VSVG) protein, thus it entered the cells without a specific recognition of any receptor. Both the viruses carried the Green Fluorescent Protein (GFP) gene in their RNA genome, which was expressed and easily detected in the cells upon infection. PPE was used at the safe dose of 0.04 mg/mL, as determined by the cytotoxicity MTT assay (data not shown). As shown in **Figure 3**, when the cells were infected by the lentivirus carrying the S protein in the presence of PPE, the percentage of GFP fluorescent cells (infected cells) was almost significantly abolished after 72 h. Contrarily, when the cells were infected by the lentivirus carrying VSVG protein, the percentage of infected cells was reduced only by 18%, suggesting a specific inhibitory effect of PPE towards Spike/ACE2 binding.

**Figure 3:**
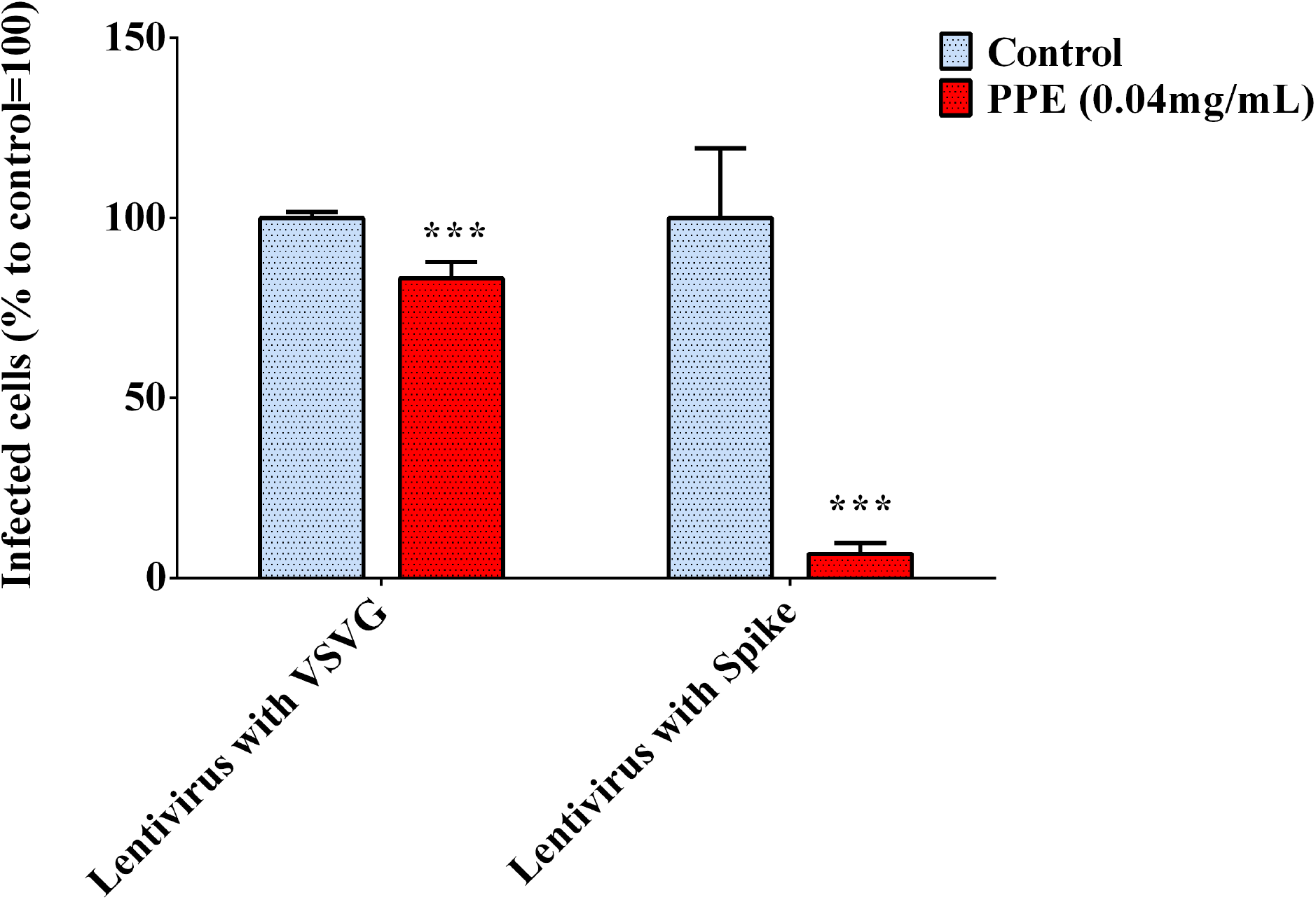
Infection rate of Spike SarsCov2 pseudo-typed lentivirus in human renal cells (HK2), determined by GFP fluorescence measure. The results are the averages of six independent experiments, expressed as percentages respect to control arbitrarily set as 100%. The error bars represent standard deviations and the asterisks indicate statistically significant values (*** p value is between 0.0001 to 0.001) according to T test.

To investigate whether PPE could regulate host genes involved in the virus uptake, we measured the expression level of ACE2 and TMPRSS2 genes in HK2 cells treated with the extract for 72 h. As reported in **Figure 4**, the gene expression analysis showed that the treatment of HK2 cells with the PPE at 0.04 mg/mL reduced the level of ACE2 and TMPRSS2 gene expression by 30% and 70% respectively. This suggested that PPE, besides Spike/ACE2 binding inhibition, was able to downregulate the expression of two genes responsible for the virus access into the cells.

**Figure 4:**
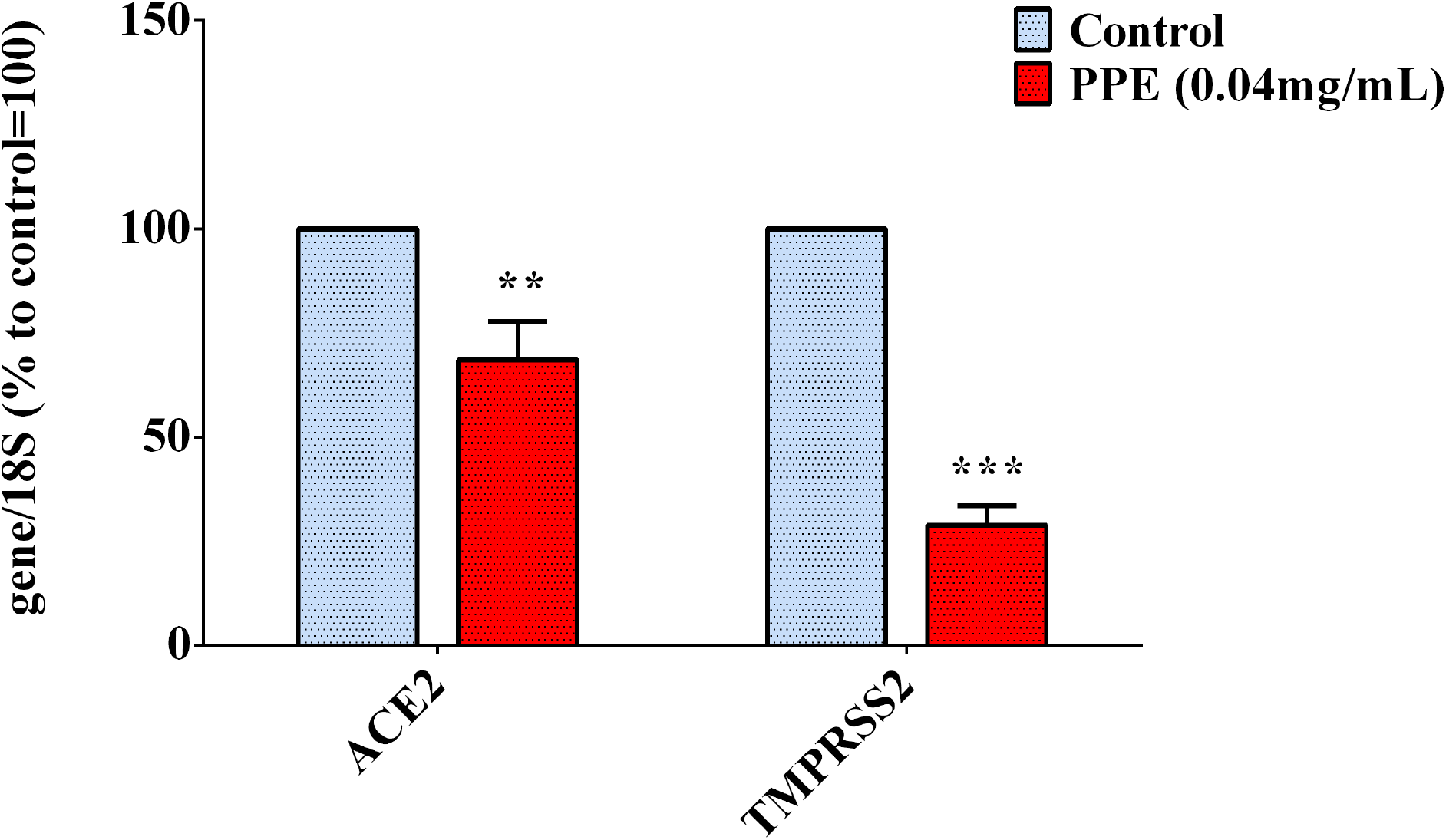
Gene expression analysis in HK2 cells treated with PPE for 72 h. The results are the averages of three independent RT-PCR experiments. The values are expressed as percentages respect to control arbitrarily set as 100%. The error bars represent standard deviations and the asterisks indicate statistically significant values (** p value is between 0.001 to 0.01; *** 0.0001 to 0.001) according to T test.

As the expression of TMPRSS2 was mainly regulated by androgens^52,53^, we analysed if PPE inhibited the 5α-Reductase activity, primary enzyme involved in DiHydroTestosterone (DHT) synthesis. As shown in **Figure 5**, PPE at 0.04mg/mL reduced the activity of the 5α-Reductase by 65% in Human Follicle Dermal Papilla cells (HFDPC), after stimulation by testosterone. This effect was similar to that obtained by finasteride, used as positive control^54^.

**Figure 5:**
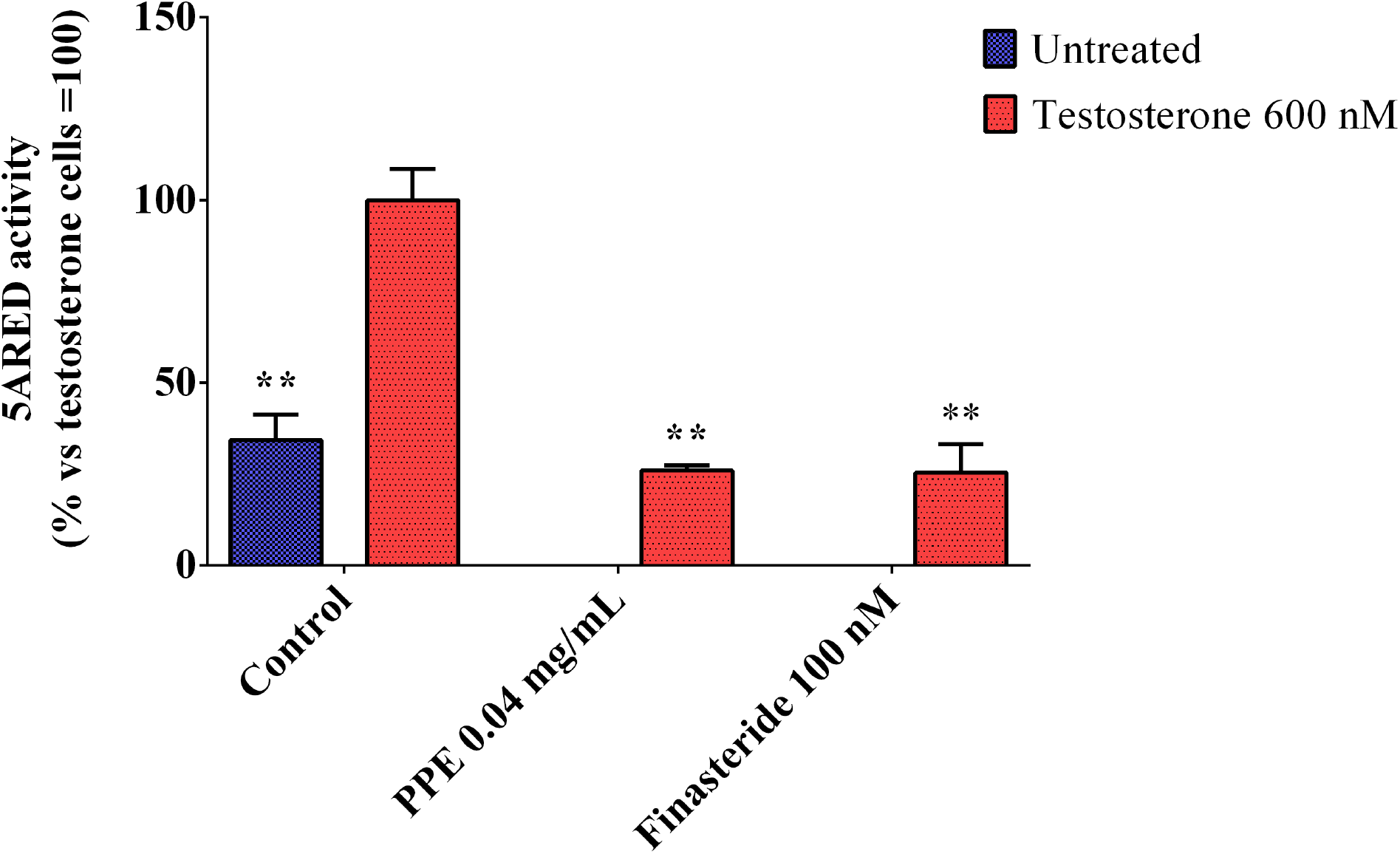
5 **α** Reductase activity in Human Follicle Dermal Papilla cells (HFDPC) stimulated with testosterone 600 nM and treated with either PPE or finasteride 100 nM. The results are the averages of three independent experiments, expressed as percentages respect to testosterone stimulated cells, arbitrarily set as 100%. The error bars represent standard deviations and the asterisks indicate statistically significant values (** p value is between 0.001 to 0.01) according to T test.

### Activity of PPE on SarsCov-2 main protease

The regulation of the 3CL protease, one of the main proteins involved in the virus replication, by the extract was investigated by incubating the enzyme with PPE and its main components, PC, EA and GA. The results, reported in the **Figure 6**, indicated that PPE, at both concentrations, inhibited the activity of the 3CL protease up to 80%. Among the compounds, PC was the most effective in inhibiting the enzymatic activity (about 50%), EA inhibited only by 10%, while GA did not have any effect, suggesting a synergic effect of the PPE polyphenols in inhibiting the protease activity.

**Figure 6:**
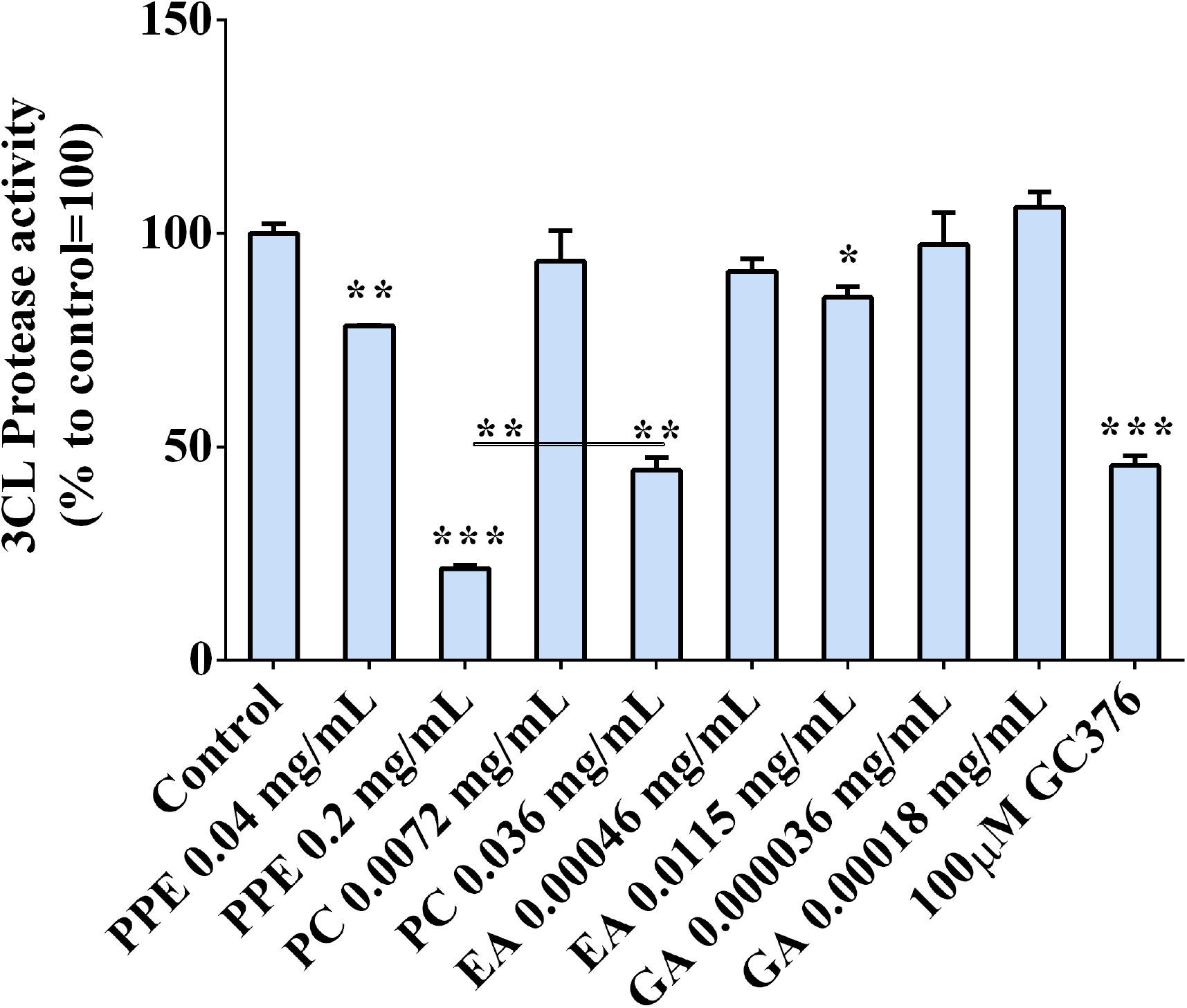
3CL protease activity in the presence of PPE, the main extract compounds (PC, EA and GA) or GC376 used as positive control. The results are the averages of three independent experiments, expressed as percentages respect to control arbitrarily set as 100%. The error bars represent standard deviations and the asterisks indicate statistically significantly values (*p value is between 0.01 to 0.05; ** 0.001 to 0.01; *** 0.0001 to 0.001) according to T test.

## Conclusions

The activity of plant secondary metabolites against SARS-CoV-2 infection and replication has been extensively reviewed in the last months^55-58^ and many studies, based on *in silico* approaches, suggested some of them as potential drug candidates for COVID-19 treatment^59^. Both viral structural proteins, like Spike, and non-structural proteins, such as 3CL^pro^, PL^pro^ and RdRp, have been proposed as valuable targets for anti-SARS-CoV-2 therapeutic strategies. Through molecular docking analyses Khalifa *et al.* 2020 found that some hydrolysable tannins, in particular pedunculagin, tercatain, and castalin, might serve as potential inhibitors of SARS‐CoV‐2 as they were able to specifically bind the 3CL protease catalytic site^60^.

In parallel studies Hariprasad *et al.* 2020 tested the virtual interaction between many plant secondary metabolites and four target proteins involved in Covid-19, the host protease TMPRSS2 and the three virus proteins, Spike, Main Protease and RNA-dependent RNA polymerase, and predicted among the class of triterpenoids the most active compounds in blocking the Spike binding site^61^. Bhatia *et al.*^37^ (2020) also identified PC among dietary polyphenols as potential inhibitor of Spike and other viral proteases. On the other side, human targets have been taken under consideration as well: ACE2 is certainly the most explored as it turned out to be the main “door lock“ that the virus used to get into the cells. However, ACE2 does have a pivotal role in many physio-pathological processes in human tissues either, thus targeting this enzyme needs careful evaluation to ensure that the benefit-risk balance turns favorable^62-64^.

In the present study, we found that the polyphenols contained in a hydro-ethanolic extract derived from pomegranate peels inhibited the interaction between Spike and ACE2 and reduced the activity of the viral 3CL protease *in vitro*, potentially suggesting the use of the extract as adjuvant in the treatment against SARS-CoV-2 infections. Data showed that the most effective polyphenols in the extract were PC and EA, possibly through a chemical interaction of the hydroxyl and galloyl groups in their molecules with the amino acid lateral groups of the Spike protein, as supported by other studies^65,66^. The inhibitory effect on Spike/ACE2 binding was confirmed by experiments with a pseudotyped lentivirus, whose entry into the human cells was dependent on Spike protein. Consistent with the *in vitro* observations, our data showed that the lentivirus infection was almost completely abolished by the polyphenol-containing PPE. This inhibition was also associated with a downregulation of the gene expression of both ACE2 and the protease TMPRSS2, the one involved in Spike priming. Moreover, we also provided evidence that PPE was able to inhibit the activity of the 3CL protease up to 80%, suggesting that PPE may have multiple biological roles in reducing the virus chance to anchor the cells and get internalized.

In conclusion, inhibiting Spike/ACE2 binding still represent one of the most popular strategies to control SARS-CoV-2, and polyphenol-rich extracts have been proposed as bioactive ingredients in pharmaceutical, nutraceutical and/or cosmetic formulations, as they represent promising candidates to reduce virus infection and replication. In agreement with our results, a recent report demonstrated that a pomegranate juice was effective in reducing the infectious capacity of Sars-Cov2 and influenza virus in VeroE6 cells suggesting an antiviral activity of both viruses^66^. The study here presented paves the way for a deeper investigation on the activity of pomegranate peel polyphenols in preventing SARS-CoV-2 infection *in vivo* and it may also promote new ideas on how reuse agroindustry byproducts for medical and health care applications.

## Abbreviations

PPE: pomegranate peel extract
S: spike protein
PC: punicalagin
EA: ellagic acid
GA: gallic acid
EAs: ellagic acid derivatives
ACE2: angiotensin-converting enzyme 2
SARS-CoV-2: Severe Acute Respiratory Syndrome Coronavirus-2
COVID-19: Coronavirus Disease 19

---

**Supplementary Figure 1:**
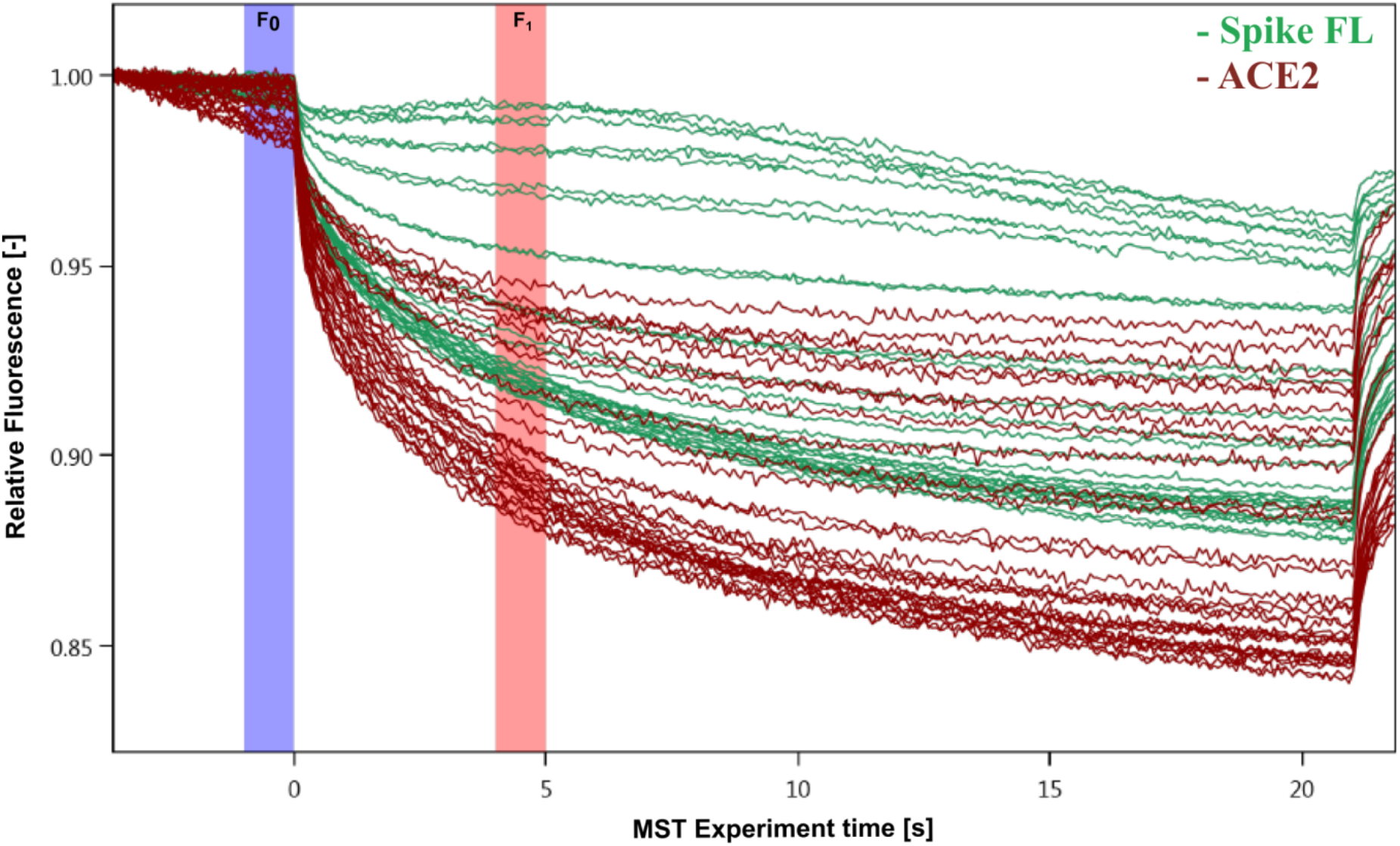
MST traces of titrations of PPE against Spike (green) and ACE2 (red); F0 and F1 correspond to the fluorescence of the unbound state and the bound state respectively.

**Supplementary Figure 2:**
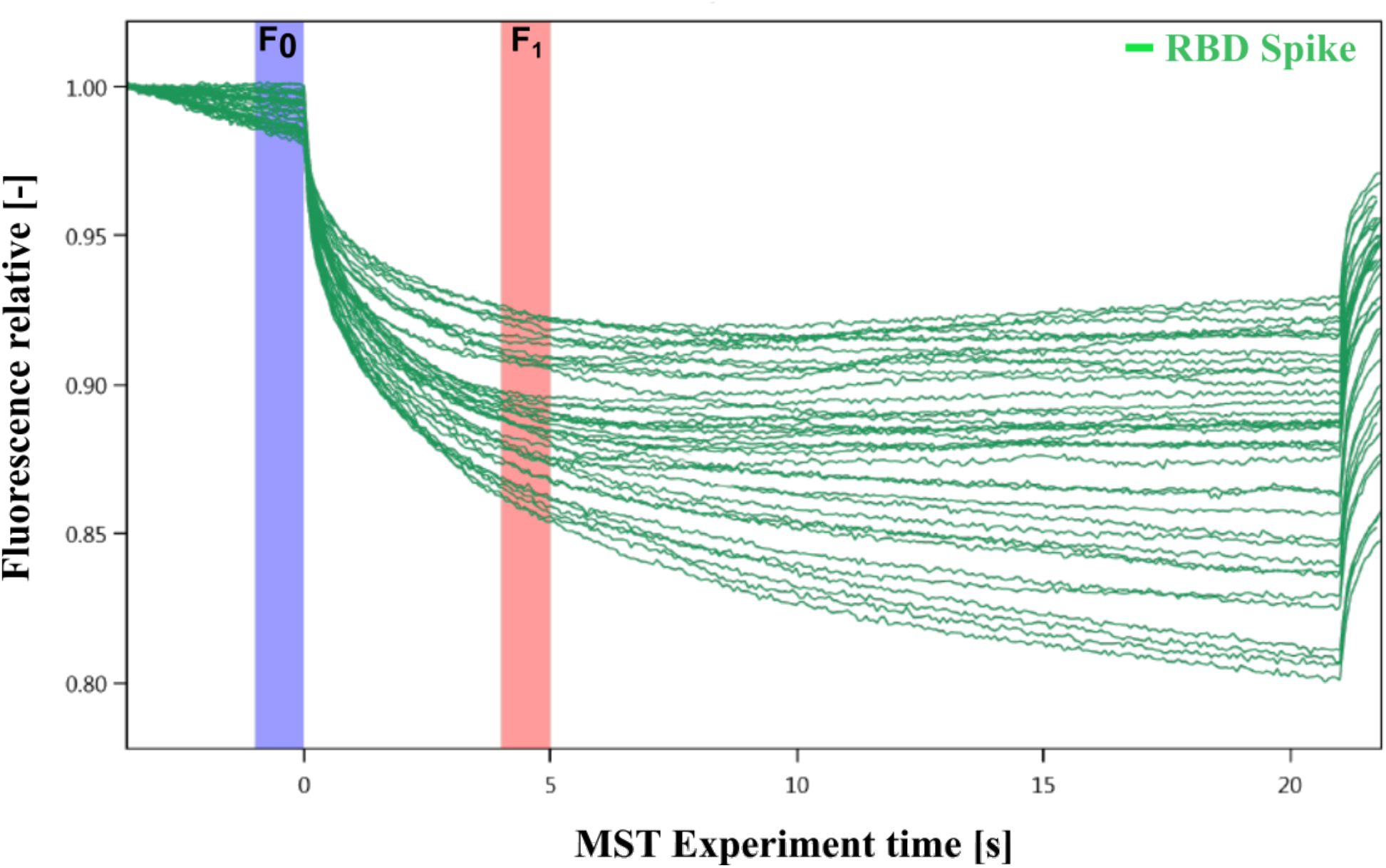
MST traces of titrations of PPE against RBD Spike; F0 and F1 correspond to the fluorescence of the unbound state and the bound state respectively.

## References

1. Isah T. Stress and defense responses in plant secondary metabolites production. Biol Res. 2019;52(1):39. Published 2019 Jul 29. doi:10.1186/s40659-019-0246-3

2. Yang L, Wen KS, Ruan X, Zhao YX, Wei F, Wang Q. Response of Plant Secondary Metabolites to Environmental Factors. Molecules. 2018;23(4):762. Published 2018 Mar 27. doi:10.3390/molecules23040762

3. Atanasov AG, Waltenberger B, Pferschy-Wenzig EM, et al. Discovery and resupply of pharmacologically active plant-derived natural products: A review. Biotechnol Adv. 2015;33(8):1582–1614. doi:10.1016/j.biotechadv.2015.08.001

4. Nasri H, Baradaran A, Shirzad H, Rafieian-Kopaei M. New concepts in nutraceuticals as alternative for pharmaceuticals. Int J Prev Med. 2014;5(12):1487–1499.

5. Barbulova A, Colucci G and Apone F. New trends in cosmetics: by-products of plant origin and their potential use as cosmetic active ingredients. Cosmetics 2015 2,82–92 doi:10.3390/cosmetics2020082

6. Ejaz Ahmed, Muhammad Arshad, Muhammad Zakriyya Khan, Muhammad Shoaib Amjad, Huma Mehreen Sadaf, Iqra Riaz, Sidra Sabir, Nabila Ahmad and Sabaoon. Secondary metabolites and their multidimensional prospective in plant life. Journal of Pharmacognosy and Phytochemistry 2017; 6(2): 205–214

7. Gorzynik-Debicka M, Przychodzen P, Cappello F, Kuban-Jankowska A, Marino Gammazza A, Knap N, Wozniak M, Gorska-Ponikowska M. Potential Health Benefits of Olive Oil and Plant Polyphenols. Int J Mol Sci. 2018 Feb 28;19(3):686. doi:10.3390/ijms19030686

8. Serreli G, Deiana M. In vivo formed metabolites of polyphenols and their biological efficacy. Food Funct. 2019 Nov 1;10(11):6999–7021. doi:10.1039/c9fo01733j. Epub 2019 Oct 29. PMID: 31659360.

9. Yiu, C.-Y.; Chen, S.-Y.; Chang, L.-K.; Chiu, Y.-F.; Lin, T.-P. Inhibitory Effects of Resveratrol on the Epstein-Barr Virus Lytic Cycle. Molecules 2010, 15, 7115–7124.

10. Zhang L, Li Y, Gu Z, et al. Resveratrol inhibits enterovirus 71 replication and pro-inflammatory cytokine secretion in rhabdosarcoma cells through blocking IKKs/NF-κB signaling pathway. PLoS One. 2015;10(2):e0116879. Published 2015 Feb 18. doi:10.1371/journal.pone.0116879

11. Annunziata G, Maisto M, Schisano C, et al. Resveratrol as a Novel Anti-Herpes Simplex Virus Nutraceutical Agent: An Overview. Viruses. 2018;10(9):473. Published 2018 Sep 3. doi:10.3390/v10090473

12. Lin CJ, Lin HJ, Chen TH, et al. Polygonum cuspidatum and its active components inhibit replication of the influenza virus through toll-like receptor 9-induced interferon beta expression [published correction appears in PLoS One. 2015;10(4):e0125288]. PLoS One. 2015;10(2):e0117602. Published 2015 Feb 6. doi:10.1371/journal.pone.0117602

13. Zang N, Xie X, Deng Y, et al. Resveratrol-mediated gamma interferon reduction prevents airway inflammation and airway hyperresponsiveness in respiratory syncytial virus-infected immunocompromised mice. J Virol. 2011;85(24):13061–13068. doi:10.1128/JVI.05869-11

14. Denaro M, Smeriglio A, Barreca D, et al. Antiviral activity of plants and their isolated bioactive compounds: An update. Phytother Res. 2020;34(4):742–768. doi:10.1002/ptr.6575

15. Lin LT, Chen TY, Chung CY, et al. Hydrolyzable tannins (chebulagic acid and punicalagin) target viral glycoprotein-glycosaminoglycan interactions to inhibit herpes simplex virus 1 entry and cell-to-cell spread. J Virol. 2011;85(9):4386–4398. doi:10.1128/JVI.01492-10

16. Nutan, Modi M, Goel T, et al. Ellagic acid & gallic acid from Lagerstroemia speciosa L. inhibit HIV-1 infection through inhibition of HIV-1 protease & reverse transcriptase activity. Indian J Med Res. 2013;137(3):540–548.

17. Zhu N, Zhang D, Wang W, et al. A Novel Coronavirus from Patients with Pneumonia in China, 2019. N Engl J Med. 2020;382(8):727–733. doi:10.1056/NEJMoa2001017

18. https://covid19.who.int/

19. Hu B, Guo H, Zhou P, et al. Characteristics of SARS-CoV-2 and COVID-19. Nat Rev Microbiol 2020. https://doi.org/10.1038/s41579-020-00459-7

20. Hoffmann M, Kleine-Weber H, Schroeder S, et al. SARS-CoV-2 Cell Entry Depends on ACE2 and TMPRSS2 and Is Blocked by a Clinically Proven Protease Inhibitor. Cell. 2020;181(2):271–280.e8. doi:10.1016/j.cell.2020.02.052

21. Remuzzi A, Remuzzi G. COVID-19 and Italy: what next?. Lancet. 2020;395(10231):1225–1228. doi:10.1016/S0140-6736(20)30627-9

22. Guan WJ, Ni ZY, Hu Y, et al. Clinical Characteristics of Coronavirus Disease 2019 in China. N Engl J Med. 2020;382(18):1708–1720. doi:10.1056/NEJMoa2002032

23. Espinosa OA, Zanetti ADS, Antunes EF, Longhi FG, Matos TA, Battaglini PF. Prevalence of comorbidities in patients and mortality cases affected by SARS-CoV2: a systematic review and meta-analysis. Rev Inst Med Trop Sao Paulo. 2020;62:e43. Published 2020 Jun 22. doi:10.1590/S1678-9946202062043

24. Onder G, Rezza G, Brusaferro S. Case-Fatality Rate and Characteristics of Patients Dying in Relation to COVID-19 in Italy [published correction appears in JAMA. 2020 Apr 28;323(16):1619]. JAMA. 2020;323(18):1775–1776. doi:10.1001/jama.2020.4683

25. Pirone L, Del Gatto A, Di Gaetano S, et al. A Multi-Targeting Approach to Fight SARS-CoV-2 Attachment. Front Mol Biosci. 2020;7:186. Published 2020 Aug 3. doi:10.3389/fmolb.2020.00186

26. Tresoldi I, Sangiuolo CF, Manzari V, Modesti A. SARS-COV-2 and infectivity: Possible increase in infectivity associated to integrin motif expression [published online ahead of print, 2020 Apr 4]. J Med Virol. 2020;10.1002/jmv.25831. doi:10.1002/jmv.25831

27. Davidson AM, Wysocki J, Batlle D. Interaction of SARS-CoV-2 and Other Coronavirus With ACE (Angiotensin-Converting Enzyme)-2 as Their Main Receptor: Therapeutic Implications. Hypertension. 2020;76(5):1339–1349. doi:10.1161/HYPERTENSIONAHA.120.15256

28. Anand K, Ziebuhr J, Wadhwani P, Mesters JR, Hilgenfeld R. Coronavirus main proteinase (3CLpro) structure: basis for design of anti-SARS drugs. Science. 2003;300(5626):1763–1767. doi:10.1126/science.1085658

29. Shin D, Mukherjee R, Grewe D, et al. Papain-like protease regulates SARS-CoV-2 viral spread and innate immunity [published online ahead of print, 2020 Jul 29]. Nature. 2020;10.1038/s41586-020-2601-5. doi:10.1038/s41586-020-2601-5

30. Ahmad J, Ikram S, Ahmad F, Rehman IU, Mushtaq M. SARS-CoV-2 RNA Dependent RNA polymerase (RdRp) - A drug repurposing study. Heliyon. 2020;6(7):e04502. Published 2020 Jul 23. doi:10.1016/j.heliyon.2020.e04502

31. Zhang Y, Geng X, Tan Y, et al. New understanding of the damage of SARS-CoV-2 infection outside the respiratory system. Biomed Pharmacother. 2020; 127:110195. doi:10.1016/j.biopha.2020.110195

32. Pascarella G, Strumia A, Piliego C, et al. COVID-19 diagnosis and management: a comprehensive review. J Intern Med. 2020;288(2):192–206. doi:10.1111/joim.13091

33. https://biorender.com/covid-vaccine-tracker

34. Dube T, Ghosh A, Mishra J, Kompella UB, Panda JJ. Repurposed Drugs, Molecular Vaccines, Immune-Modulators, and Nanotherapeutics to Treat and Prevent COVID-19 Associated with SARS-CoV-2, a Deadly Nanovector [published online ahead of print, 2020 Oct 25]. Adv Ther (Weinh). 2020;2000172. doi:10.1002/adtp.202000172

35. Khaerunnisa S, Kurniawan H, Awaluddin R, Suhartati S, Soetjipto S. Potential Inhibitor of COVID-19 Main Protease (M<sup>pro</sup>) From Several Medicinal Plant Compounds by Molecular Docking Study. Preprints.org; 2020. DOI:10.20944/preprints202003.0226.v1.

36. Adem S, Eyupoglu V, Sarfraz I, Rasul A, Ali M. Identification of Potent COVID-19 Main Protease (Mpro) Inhibitors from Natural Polyphenols: An in Silico Strategy Unveils a Hope against CORONA. Preprints 2020, 2020030333 doi:10.20944/preprints202003.0333.v1.

37. Bhatia S, Giri S, Lal AF, Singh S. Identification of potential inhibitors of dietary polyphenols for SARS-CoV-2 M protease: An in silico study. Tropical Public Health 2020; 1(1): 21–29

38. Reddy MK, Gupta SK, Jacob MR, Khan SI, Ferreira D. Antioxidant, antimalarial and antimicrobial activities of tannin-rich fractions, ellagitannins and phenolic acids from Punica granatum L. Planta Med. 2007;73(5):461–467. doi:10.1055/s-2007-967167

39. Akhtar S, Ismail T, Fraternale D, Sestili P. Pomegranate peel and peel extracts: chemistry and food features. Food Chem. 2015;174:417–425. doi:10.1016/j.foodchem.2014.11.035

40. Landete J. Ellagitannins, ellagic acid and their derived metabolites: a review about source, metabolism, functions and health, Food Res. Int., 2011, 44, 1150–1160.

41. Viuda-Martos M, Fernández-López J and Pérez-Álvarez J. Pomegranate and its many functional components as related to human health: a review. Compr. Rev. Food Sci. Food Saf., 2010, 9, 635–654.

42. Howell AB, D'Souza DH. The pomegranate: effects on bacteria and viruses that influence human health. Evid Based Complement Alternat Med. 2013;2013:606212. doi:10.1155/2013/606212

43. Moradi MT, Karimi A, Shahrani M, Hashemi L, Ghaffari-Goosheh MS. Anti-Influenza Virus Activity and Phenolic Content of Pomegranate (Punica granatum L.) Peel Extract and Fractions. Avicenna J Med Biotechnol. 2019;11(4):285–291.

44. Houston DMJ, Bugert JJ, Denyer SP, Heard CM. Potentiated virucidal activity of pomegranate rind extract (PRE) and punicalagin against Herpes simplex virus (HSV) when co-administered with zinc (II) ions, and antiviral activity of PRE against HSV and aciclovir-resistant HSV [published correction appears in PLoS One. 2017 Nov 20;12 (11):e0188609]. PLoS One. 2017;12(6):e0179291. Published 2017 Jun 30. doi:10.1371/journal.pone.0179291

45. Malviya S, Arvind, Jha A, Hettiarachchy N. Antioxidant and antibacterial potential of pomegranate peel extracts. J Food Sci Technol. 2014;51(12):4132–4137. doi:10.1007/s13197-013-0956-4.

46. Colantuono A, Vitaglione P, Ferracane R, Campanella OH, Hamaker BR. Development and functional characterization of new antioxidant dietary fibers from pomegranate, olive and artichoke by-products. Food Res Int. 2017;101:155–164. doi:10.1016/j.foodres.2017.09.001

47. Re R, Pellegrini N, Proteggente A, Pannala A, Yang M, Rice-Evans C. Antioxidant activity applying an improved ABTS radical cation decolorization assay. Free Radic Biol Med. 1999;26(9-10):1231–1237. doi:10.1016/s0891-5849(98)00315-3

48. Fischer UA, Carle R, Kammerer DR. Identification and quantification of phenolic compounds from pomegranate (Punica granatum L.) peel, mesocarp, aril and differently produced juices by HPLC-DAD-ESI/MS(n). Food Chem. 2011;127(2):807–821. doi:10.1016/j.foodchem.2010.12.156

49. Lu J, Ding K, & Yuan Q. Determination of punicalagin isomers in pomegranate husk. Chromatographia, 2008 68, 303–306.

50. Marchi LB, Monteiro AR, Mikcha J, Santos A, Chinelatto M, Marques D, Dacome A and Costa SC. Evaluation of antioxidant and antimicrobial capacity of pomegranate peel extract (*Punica granatum*l.) Under different drying temperatures. Chemical engineering transactions, 2015 44, 121–126.

51. Koka V, Huang XR, Chung AC, Wang W, Truong LD, Lan HY. Angiotensin II up-regulates angiotensin I-converting enzyme (ACE), but down-regulates ACE2 via the AT1-ERK/p38 MAP kinase pathway. Am J Pathol. 2008;172(5):1174–1183. doi:10.2353/ajpath.2008.070762

52. Oyelowo O, Okafor C, Ajibare A, et al. Fatty Acids in Some Cooking Oils as Agents of Hormonal Manipulation in a Rat Model of Benign Prostate Cancer. Niger J Physiol Sci. 2019;34(1):69–75. Published 2019 Jun 30.

53. Hong MY, Seeram NP, Heber D. Pomegranate polyphenols down-regulate expression of androgen-synthesizing genes in human prostate cancer cells overexpressing the androgen receptor. J Nutr Biochem. 2008;19(12):848–855. doi:10.1016/j.jnutbio.2007.11.006

54. Rattanachitthawat N, Pinkhien T, Opanasopit P, Ngawhirunpat T, Chanvorachote P. Finasteride Enhances Stem Cell Signals of Human Dermal Papilla Cells. In Vivo. 2019;33(4):1209–1220. doi:10.21873/invivo.11592

55. Antonio AdS, Wiedemann LSM, and Veiga-Junior VF. Natural products' role against COVID-19. RSC Adv., 2020,10, 23379–23393

56. Sayed AM, Khattab AR, AboulMagd AM, Hassan HM, Rateb ME, Zaid H, Abdelmohsen UR Nature as a treasure trove of potential anti-SARS-CoV drug leads: a structural/mechanistic rationale RSC Adv., 2020, 10, 19790–19802

57. Weng JK. Plant Solutions for the COVID-19 Pandemic and Beyond: Historical Reflections and Future Perspectives. Mol Plant. 2020;13(6):803–807. doi:10.1016/j.molp.2020.05.014

58. Singh YD, Jena B, Ningthoujam R, et al. Potential bioactive molecules from natural products to combat against coronavirus. ADV TRADIT MED (ADTM) 2020. https://doi.org/10.1007/s13596-020-00496-w

59. Majumder R, Mandal M. Screening of plant-based natural compounds as a potential COVID-19 main protease inhibitor: an in silicodocking and molecular dynamics simulation approach [published online ahead of print, 2020 Sep 8]. J Biomol Struct Dyn. 2020;1–16. doi:10.1080/07391102.2020.1817787

60. Khalifa I, Zhu W, Mohammed HHH, Dutta K, Li C. Tannins inhibit SARS-CoV-2 through binding with catalytic dyad residues of 3CLpro: An in silico approach with 19 structural different hydrolysable tannins [published online ahead of print, 2020 Aug 11]. J Food Biochem. 2020;e13432. doi:10.1111/jfbc.13432

61. Hariprasad P, Gowtham HG, Monu DO, Ajay Y, Gourav C, Vasantharaja R, Bhani K, Koushalya S, Shazia S, Priyanka G, Faiz A, Leena C. Exploring structurally diverse plant secondary metabolites as a potential source of drug targeting different molecular mechanisms of Severe Acute Respiratory Syndrome Coronavirus-2 (SARS-CoV-2) pathogenesis: An in silico approach. 10.21203/rs.3.rs-27313/v1

62. Mostafa-Hedeab G. ACE2 as Drug Target of COVID-19 Virus Treatment, Simplified Updated Review. Rep Biochem Mol Biol. 2020;9(1):97–105. doi:10.29252/rbmb.9.1.97

63. Lacroix C, de Wouters T, Chassard C. Integrated multi-scale strategies to investigate nutritional compounds and their effect on the gut microbiota. Curr Opin Biotechnol. 2015;32:149–155. doi:10.1016/j.copbio.2014.12.009

64. Khare P, Sahu U, Pandey SC, Samant M. Current approaches for target-specific drug discovery using natural compounds against SARS-CoV-2 infection. Virus Res. 2020;290:198169. doi:10.1016/j.virusres.2020.198169

65. Maiti S, BanerjeeA. Epigallocatechin gallate and theaflavin gallate interaction in SARS‐CoV‐2 spike‐protein central channel with reference to the hydroxychloroquine interaction: Bioinformatics and molecular docking study. Drug Dev Res. 2020; 1–11. https://doi.org/10.1002/ddr.21730

66. Conzelmann C, Weil T, Groß R, Jungke P, Frank B, Eggers M, Müller JA, Münch J. Antiviral activity of plant juices and green tea against SARS-CoV-2 and influenza virus *in vitro*. bioRxiv 2020.10.30.360545; doi:https://doi.org/10.1101/2020.10.30.360545

